# Magnetophospenes in humans exposed to ELF MF up to 50 mT, a threshold study

**DOI:** 10.1101/439968

**Authors:** A Legros, J Modolo, M Corbacio, D Goulet, M Plante, M Souques, F Deschamps, G Ostiguy, J Lambrozo, AW Thomas

## Abstract

Although magnetophosphene perception is the most reliable reported effect on acute human neurophysiological responses to extremely low frequency (ELF) magnetic field (MF) exposure, current knowledge is based on small sample size, non-replicated experiments. In this study, we established MF levels triggering magnetophosphenes at 20, 50, 60 and 100 Hz in humans. Magnetophosphene perception and EEG were collected in 55 magnetic flux density conditions randomly delivered in each frequency group (2 experiments, total n=145). Results indicate that threshold values 1) need to be reported as a function of dB/dt instead of flux density, and 2) are frequency-dependent (higher sensitivity to lower frequencies). No clear trend was found in EEG data.

## INTRODUCTION

In the Extremely Low Frequency (ELF) range (< 300 Hz), the threshold for magnetophosphene perception constitutes the experimental basis of international recommendations regarding exposure to magnetic fields (MF) [1, 2]. Both the International Commission for Non-Ionizing Radiation Protection (ICNIRP) and the Institute of Electrical and Electronics Engineers (IEEE) provide international recommendations regarding human exposure to ELF MF based on the threshold for magnetophosphene perception as reported by Lövsund in the early eighties [3]. However, significant uncertainties persist regarding this threshold [4, 5]. Magnetophosphene perception is reported to be optimal at 20 Hz (threshold between 5 and 10 mT) and depreciated at both higher and lower frequencies [6, 7]. However, this conflicts with recent studies using transcranial electrical stimulation reporting improved phosphene perception between 8 and 15 Hz in dark conditions [8, 9]. Also, only extrapolated data are available at 60 Hz, and no human experimental data is available to date. The aim of this project was therefore to establish in humans the thresholds for acute, objective and quantifiable responses in humans exposed to ELF MF of up to 50 mT, and more specifically at 20/50/60/100 Hz. Since magnetophosphenes are by definition a flickering light visual perception, we hypothesized that magnetophosphene perception is associated with a decrease in EEG alpha activity (8-12 Hz) in the visual/occipital cortex. Reported magnetophosphene perception and associated electroencephalography (EEG) results are presented.

## MATERIAL AND METHODS

Two experiments were conducted using the exact same protocol. Experiment 1 tested only 2 frequency groups (n=30 at 50 Hz, n=34 at 60 Hz) using a prototype version of the exposure system, and experiment 2 was extended to 4 frequency groups (new volunteers, n=20 at 20, 60 and 100 Hz; n=21 at 50 Hz). Each healthy volunteer was tested in two localized exposure conditions (eyeball and occipital cortex, using a small coil) and in one global head exposure condition (see Figure 1 for an illustration of the global head exposure system). Each frequency group was scanned with 11 magnetic flux density conditions (from 0 to 50 mT_rms_, 5 mT increments) lasting five seconds each. Each magnetic flux density condition was repeated 5 times and separated with five seconds free from MF exposure. A computer program (developed in LabVIEW, National Instruments, USA) randomly assigned the order of presentation of MF flux density conditions. During this protocol, participants were sitting eyes closed in a dark room, and were asked to report magnetophosphene perception by button-press, while occipital EEG activity (O2, O1 and OZ electrodes) was continuously recorded. Each experimental condition started after 5 minutes of adaptation to darkness. An MRI-compatible 64-channel EEG system/cap/cable (Neuroscan-Compumedics Inc., Melbourne, Australia) was used, allowing EEG recording during 50 mT MF exposure without saturating EEG amplifiers (validated using EEG phantom recordings and analyses). This protocol was approved by the Health Sciences Research Ethics Board of Western University (HSREB #18882).

**Figure 1:**
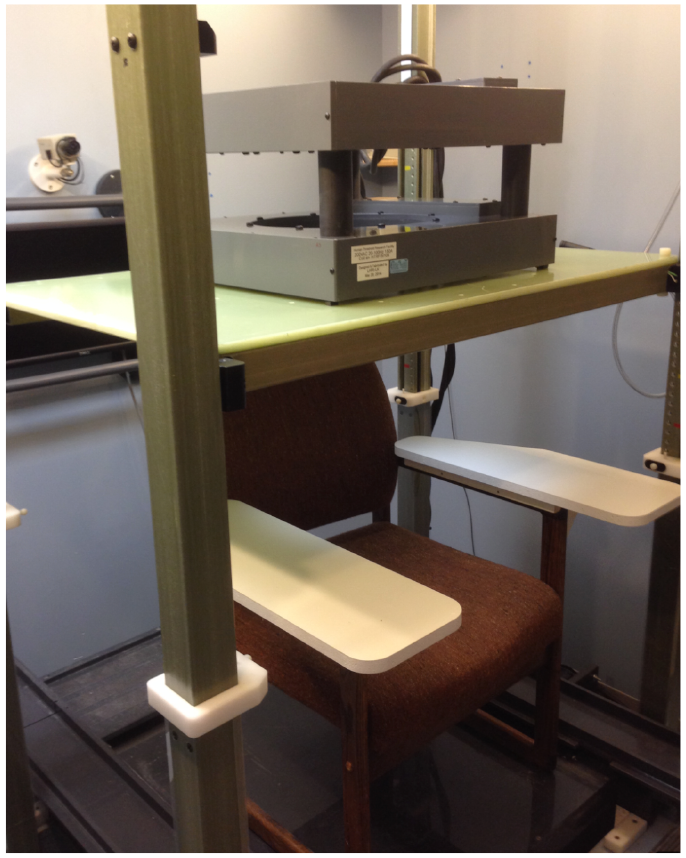
The global head exposure was delivered through a set of 2 hollow copper wire coils (enabling cool water circulation, preventing heating) of 21.4 cm diameters, vertically spaced by 20 cm. An MRI gradient amplifier powers each coil (up to 200 A capability) at the required frequency. The system is Medical Grade approved by the Canadian Standard Association (CSA).

## RESULTS

Experiments 1 and 2 group results for magnetophosphene perception in the global exposure condition are reported in this abstract. Only descriptive statistics are reported for brevity, and detailed results will be shared at the conference. In a first round of data analysis, it was decided to analyze the button-press data (i.e. selfreported magnetophosphene perceptions) as a function of the 11 flux density conditions (0 to 50 mT_rms_, 5 mT increments). Corresponding results are shown in Figure 2. Perception thresholds were found to be 10 and 15 mT at 50 and 60 Hz, respectively; and the perception rate was higher at 50 Hz as compared to 60 Hz. However, experiment 2 failed to replicate this frequency-response pattern expected from the Lösvund study (lower threshold and perception rate for lower frequencies - [3]): 1. perception thresholds were not different at 50 and 60 Hz (20 mT), and 2. although the threshold was also 20 mT at 20 Hz, the perception rate was consistently lower in this frequency group as compared to the 50 and 60 Hz groups.

**Figure 2:**
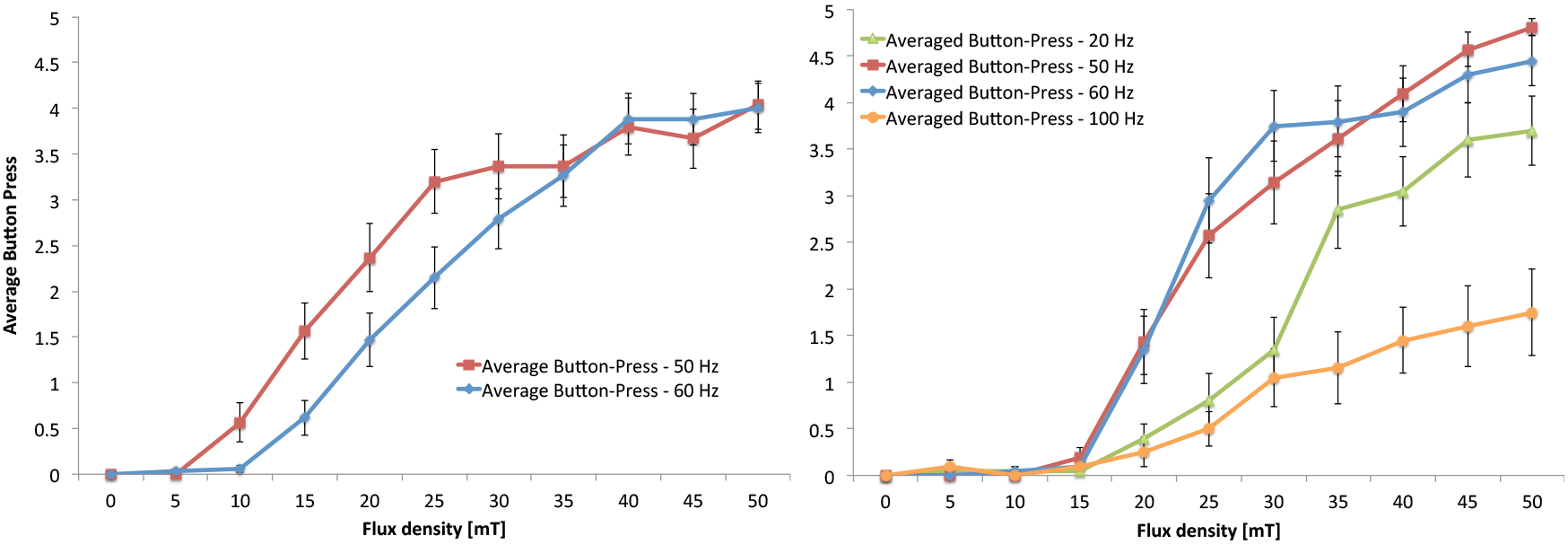
<u>*Left*</u>: Averaged number of magnetophosphene detections by button-press as a function of MF flux density for experiment 1, in the global head exposure condition. The threshold is at 10 mT in the 50 Hz group (n=30) and at 15 mT in the 60 Hz group (n=34). <u>*Right*</u>: Averaged number of magnetophosphene detections by button-press as a function of MF flux density for experiment 2, in the global head exposure condition. The threshold is at 20 mT in the 4 frequency groups, with higher sensitivity at 50 and 60 Hz (n=21 and 20 respectively) as compared to 20 and 100 Hz (n=20 in each group).

Based on these observations, it was decided to conduct a second run of analyses in both experiments, comparing the button-press data as a function of calculated dB/dt values, which are representative of the actual *in-situ* electric field at the retinal level. These results are reported in Figure 3.

**Figure 3:**
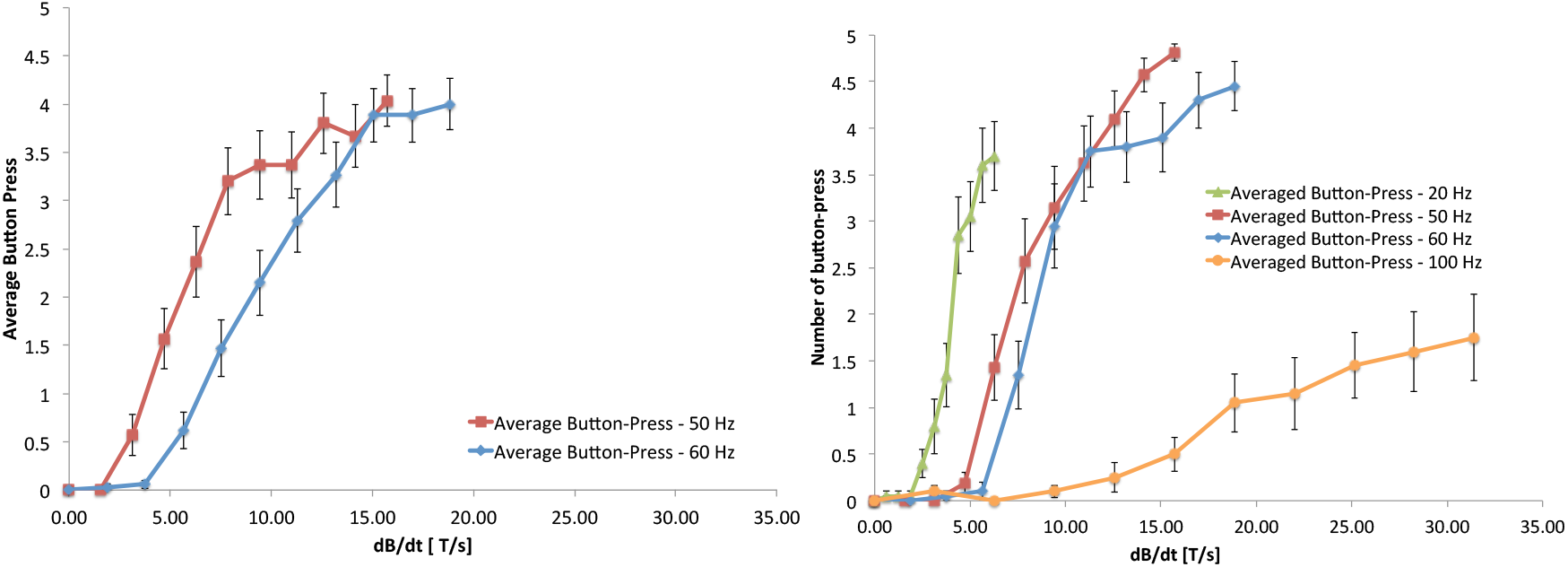
<u>*Left*</u>: Averaged number of magnetophosphene detections by button-press as a function of dB/dt for experiment 1, in the global head exposure condition. The threshold is 10 mT in the 50 Hz group (n=30) and 15 mT in the 60 Hz group (n=34). <u>*Right*</u>: Averaged number of magnetophosphene detections by button-press as a function of dB/dt for experiment 2, in the global head exposure condition. The threshold is now frequency specific: 2.51 T/s at 20 Hz, 6.28 T/s at 50 Hz, 7.54 T/s at 60 Hz, and 12.57 T/s at 100 Hz.

Results from the second round of analyses showed a frequency-dependent threshold: 2.51 T/s at 20 Hz, 6.28 T/s at 50 Hz, 7.54 T/s at 60 Hz, and 12.57 T/s at 100 Hz. In addition, the slope of the perception rate as a function of dB/dt is steeper for the lowest frequency condition, and gradually increases for higher frequencies.

The local/retinal exposure condition showed similar results than the global exposure condition (with slightly different threshold values). The local/occipital exposure condition failed to show a threshold for magnetophosphene perception.

Corresponding EEG signals from occipital electrodes O2, O1 and OZ were analyzed. Except for the global exposure condition in the experiment 1 only, no effects of the exposure conditions were found (none in experiment 2). The results will be reported at the conference.

## DISCUSSION/CONCLUSIONS

This project allowed the experimental testing of ELF MF exposures of up to 50 mT in humans at 20/50/60 and 100 Hz. These experiments concomitantly investigated a self-reported perception (magnetophosphenes) and an objective neurophysiological outcome (EEG), both acquired during MF exposure. When looking at the results reported as a function of flux density, the differential frequency-response as reported by Lövsund [3] is not replicated. It is also in contradiction with recent studies using transcranial electrical stimulation directly applied with electrodes, which report the lowest perception threshold between 8 and 15 Hz in dark conditions, and a higher threshold as the frequency increases [8, 9]. This inconsistency motivated a second data analysis strategy focusing on dB/dt values, which are linearly correlated with the actual electric field at the retinal level: choosing dB/dt as the proper metric to analyze the data allows to properly apprehend the *in-situ* electric field value, we highlighted a frequency-dependent response, with the threshold dB/dt value increasing linearly with frequency (lower threshold at lower frequencies). In addition, perception dynamics follow frequency-specific slopes, confirming greater magnetophosphene sensitivity at lower frequencies for a similar *in-situ* induced electric field level.

The absence of effect in the occipital exposure condition, as opposed to the efficacy of a single eye exposure to generate magnetophosphenes, confirm that the interaction site is in the retina and not in the occipital cortex, as it was still under consideration [10], supporting findings from Laakso and Hirata [11]. This project will provide solid experimental data acquired in humans to refine exposure guidelines, which may also offer opportunities for translational research.

In order to move forward in understanding the MF exposure frequency-dependence in magnetophosphene perception, future experimental protocols will need to be designed as a function of dB/dt values and explore higher exposure levels in order to reach a plateau in the perception rate. In addition, while our results illustrate a linear relationship between the magnetophosphene perception threshold and dB/dt in the 20-100 Hz frequency range, it will be required to investigate the full ELF spectrum (up to 300 Hz) in order to fully document the frequency response curve and improve our understanding of underlying mechanisms. It will also be important to verify magnetophosphene perception is actually facilitated at 10-15 Hz, as reported by electrophosphene studies. These experimental confirmations would support the hypothesis that retinal rods are the main origin of magnetophosphene perception.

## ACKNOWLEDGEMENTS

This work is supported by Hydro-Québec, Réseau de Transport d’Électricité, Électricité de France, NationalGrid, Energy Network Association (ENA), the Electric Power Research Institute (EPRI) and the Canadian Institutes of Health Research. The authors would like to thank Lynn Keenliside for having designed and built the exposure system.

